# Age-related deficits in rapid visuomotor decision-making

**DOI:** 10.1101/2021.02.07.430152

**Authors:** Ana Gómez-Granados, Deborah A. Barany, Margaret Schrayer, Isaac Kurtzer, Cédrick Bonnet, Tarkeshwar Singh

**Affiliations:** Department of Kinesiology, University of Georgia, Athens, GA-30602; Augusta University/University of Georgia Medical Partnership, Athens, GA-30602; Department of Biomedical Science, College of Osteopathic Medicine, New York Institute of Technology, Old Westbury, New York, NY-11568; Univ. Lille, CNRS, UMR 9193 – SCALab – Sciences Cognitives et Sciences Affectives, F-59000 Lille, France; Department of Kinesiology, The Pennsylvania State University, University Park, PA-16802

**Keywords:** Older adults, reaching, interception, dorsal-ventral interactions, object recognition

## Abstract

Many goal-directed actions that require rapid visuomotor planning and perceptual decision-making are affected in older adults, causing difficulties in execution of many functional activities of daily living. Visuomotor planning and perceptual decision-making are mediated by the dorsal and ventral visual streams, respectively, but it is unclear how age-induced changes in sensory processing in these streams contribute to declines in goal-directed actions. Previously, we have shown that in healthy adults task demands affect the integration of sensory information between the two streams and more motorically demanding tasks induce earlier decisions and more decision errors. Here, we asked the question if older adults would exhibit larger declines in interactions between the two streams during demanding motor tasks. Older adults (n=15) and young controls (n=26) performed a simple reaching task and a more demanding interception task towards virtual objects. In some blocks of trials, participants also had to select an appropriate movement based on the shape of the object. Our results showed that older adults made a similar number of initial decision errors during both the reaching and interception tasks but corrected fewer of those errors during movement. During the more demanding interception decision task, older adults made more decision- and execution-related errors than young adults, which were related to early initiation of their movements. Together, these results suggest that older adults have a reduced ability to integrate new perceptual information to guide online action, which may reflect impaired ventral-dorsal stream interactions.

**Highlights:**

- Older adults showed reduced performance in a visuomotor decision-making task
- Initial decision errors were similar between young and older adults
- Older adults were less likely to correct initial decision errors
- More demanding movements were associated with earlier and less accurate decisions

## 1. Introduction

Older adults exhibit functional deficits in many activities of daily living that require integration of sensory, cognitive, and motor processes. For example, driving requires rapid visuomotor integration to choose an appropriate motor response (e.g., judging change in traffic lights to accelerate or brake). Age-related declines in vision, decision-making, or motor control are associated with deficits in many activities of daily living (Anstey et al., 2005; McGwin Jr and Brown, 1999). These declines have been extensively investigated in isolation, however the underlying interactions that contribute to these deficits remain an open question.

Slower response times of older adults during perceptual decision-making tasks are linked to declines in both sensory processing and cognition (Dully et al., 2018). Older adults show deficits associated with sensory processing in the dorsal visual stream (Goodale and Milner, 1992), such as visual processing speed, speed discrimination, and motion perception (Biehl et al., 2017; Norman et al., 2003; Owsley, 2011). In contrast, other aspects of sensory processing that are mediated by the ventral visual stream, such as contrast discrimination (Delahunt et al., 2008; McGovern et al., 2018; Tulunay-Keesey et al., 1988) and color perception (Delahunt et al., 2005; Elliott et al., 2007; Jung and Kline, 2010) remain relatively intact. Overall, compared to the ventral stream, the dorsal stream exhibits early age-related decline in older adults (Langrová, J et al., 2006; Sciberras-Lim and Lambert, 2017). These differential age-related rates of decline in the two streams could affect many activities of daily living that require continuous and time-sensitive interactions between the two streams (Barany et al., 2020), but this has remained unexamined.

We hypothesized that older adults would exhibit impaired interactions between the dorsal and ventral stream during rapid visuomotor decision-making. To engage the ventral stream, we asked participants to select one of two alternative movements based on their judgment of the target’s shape. We predicted that older adults would make more decision errors and be less likely to make appropriate movement adjustments during decision-making. Furthermore, we used manual reaching and interception versions of the task to engage the dorsal visual stream differentially. Relative to reaching movements, planning of interception movements is more challenging as it also requires motion estimation between the moving object and the body (Brenner and Smeets, 2011; Merchant and Georgopoulos, 2006; Zago et al., 2009). Therefore, our second prediction was that older adults would make more decision- and execution-related errors in the interception task than the reaching task.

## 2. Methods

### 2.1. Participants

Twenty-six younger participants (16 women; 23.7 ± 5.5 years), and fifteen older participants (11 women; 69.2 ± 4.0 years) completed the study. All participants were right-handed, had no known history of neurological disorders, and had normal or corrected-to-normal vision. All the participants provided written informed consent prior to participating and were compensated for their time.Experimental procedures were approved by the Institutional Review Board at the University of Georgia.

### 2.2. Apparatus

Participants were seated on a chair while their right hand grasped the handle of a robotic manipulandum that moved in a horizontal plane (KINARM End-Point Lab, KINARM, Kingston, Ontario, Canada). Visual stimuli (including the handle location) were projected at 60 Hz from a monitor above the workspace onto a semi-transparent mirror, which occluded direct vision of the hand (Fig. 1A). The monitor displayed targets and a cursor representing the location of the right hand in a veridical horizontal plane. During the performance of the trials, the robot applied a constant background force (−3 N in the Y direction) to the handle and recorded movement position and velocity at 1000 Hz (Barany et al., 2020; Singh et al., 2017). Eye-tracking data was also recorded at 500 Hz using a video-based remote system (Eyelink 1000, SR Research, Ottawa, ON, Canada) and used to track fixations to begin each trial (see *Experimental design and procedure*), but not analyzed further for the current study.

**Figure 1:**
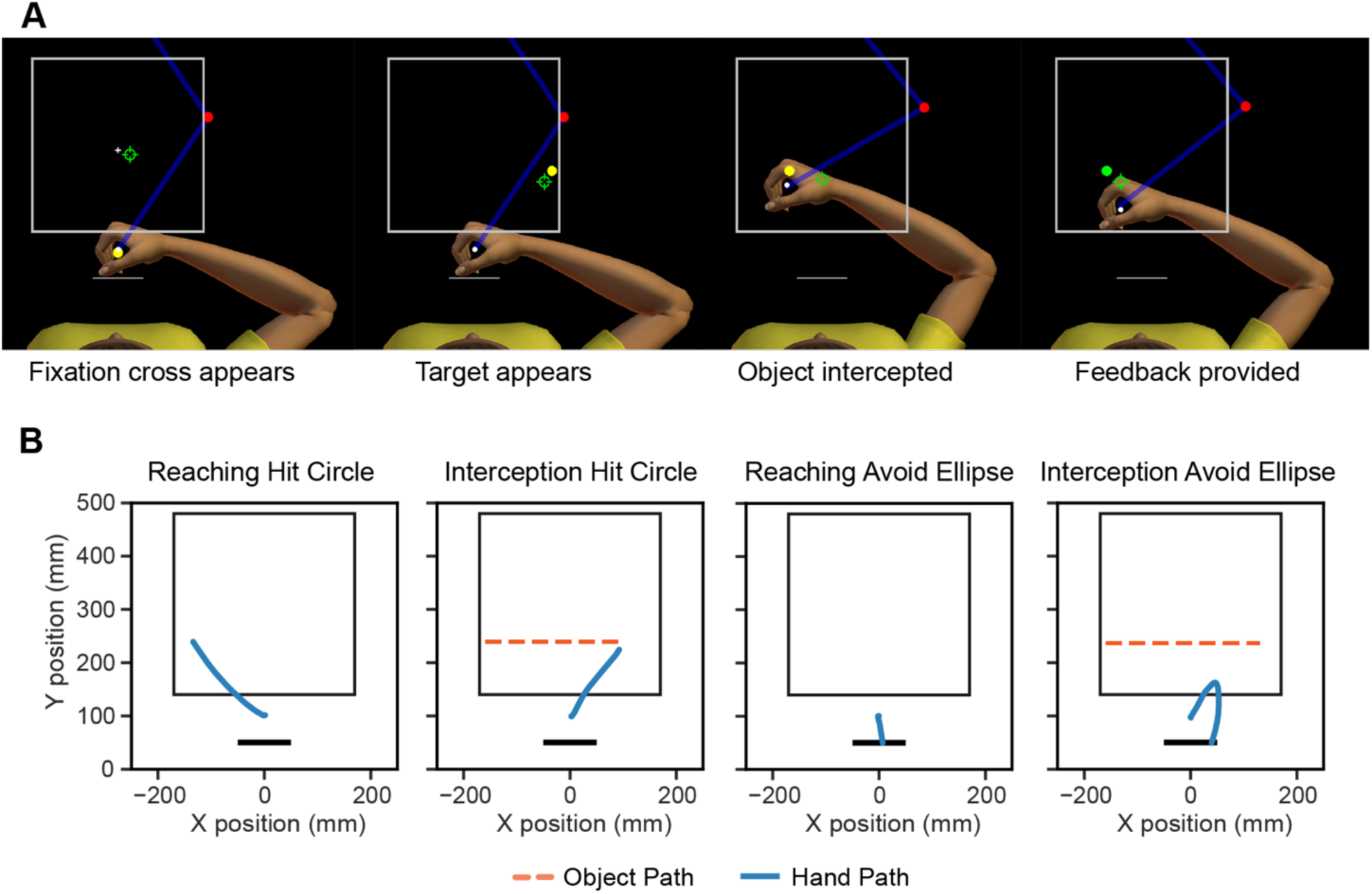
The experimental setup and an exemplar trial. A: Sample trial for interception task. The green crosshair represents the participant’s gaze location, the white cursor represents the participant’s hand location. After 500 ms of fixation on the fixation cross, a yellow target would appear on the left or right side of the screen and move towards the other end of the white box. Participants were provided with feedback when the target was intercepted or if the trial timed out. The target would turn red if the trial was unsuccessful or green if it was successful. B: Example of hand paths and different trial scenarios from a representative participant.

### 2.3. Experimental design and procedure

Experimental procedures and the young adult dataset were described in a recent study (Barany et al., 2020). In brief, participants performed rapid reaching and interception movements with their right hand (see Fig. 1A). At the beginning of each trial, participants were instructed to move their hand into a yellow circle that appeared at the starting position at the midline of the visual display. After reaching the starting position, participants were required to fixate at a fixation cross also positioned along the midline 22 cm away from the start position of the hand. After 500 ms of maintaining eye fixation (as determined from the eye-tracker) and hand position, both the fixation cross and start position disappeared. After a 200 ms delay, a yellow object was presented inside a white rectangular box either on the left or right side, ±16 cm along x-axis (see Fig. 1B). The location of the object along the y-axis could vary between 14.5-17 cm (uniform distribution). For the Reaching trials, the object stayed in its initial location. During Interception trials, the object traveled at a constant Euclidean velocity of ±40 cm/s for Fast trials and ±34 cm/s for Slow trials. The object could either be a circle (2 cm diameter) or an ellipse (minor axis = 2 cm, major axis = 1.15 × minor axis) depending on the experimental block.

In the No Decision condition, the objects were always circles for every trial in the block. In the Decision condition, the object for each trial was randomly selected to be either a circle (50% of trials) or an ellipse. For each block of trials, the object could either stay in the same position (Reaching) or move horizontally across the screen (Interception). Hence, in the No Decision blocks, participants knew beforehand that all the objects would be circles, and for the Decision blocks they were told the object could either be a circle or an ellipse. Participants were instructed to perform a reaching or interception movement as quickly and as accurately as possible when the object was a circle, and to avoid the object when it was an ellipse by moving the cursor in the opposite direction towards a bar drawn parallel to the frontal plane (Fig. 1B). For both Reaching and Interception trials, the object remained on the visual display until it was hit or the trial timed out. For the Interception trials, the maximum time on screen was determined by the object’s constant Euclidean velocity: 800 ms for Fast trials and 950 ms for Slow trials. The maximum times for the Reaching trials were also 800 ms and 950 ms, to match the Interception condition.

After the hit or the trial timed out, participants received feedback of their performance for 500 ms. The object would change the color from yellow to green (successful) or red (unsuccessful). A successful trial would be when a circle would be reached or intercepted, or when an ellipse was avoided. An unsuccessful trial would be when an ellipse was reached or intercepted, or a circle was avoided. Intertrial delay was between 1500 ms and 2000 ms.

Each participant performed 8 experimental blocks of 90 trials each (720 trials total). Blocks were randomized and consisted of a unique combination of conditions: decision type (No Decision or Decision), movement type (Reaching or Interception), and maximum trial duration (Fast or Slow). To focus our analysis on the interaction of decision-making and movement type, trials from the Fast and

Slow blocks were pooled. During Decision blocks, object shape and location of the objects along the y-axis were randomized across trials within each block. During No Decision blocks, location of the circles along the y-axis was randomized.

### 2.4. Data Analysis

All hand movement data was analyzed using MATLAB (version 9.5.0, The MathWorks, Natick, MA) and Python (version 3.7). Statistical analyses were performed in R (version 3.6.0).

Hand movement data were filtered with a fourth-order Butterworth low-pass filter with a 5 Hz cutoff (Winter, 2009). Reaction time (RT) was calculated as the time between object onset and the time when hand speed exceeded 5% of the first local peak. Trials were excluded if RT was less than 100 ms. Peak speed (PS) was calculated as the hand position’s maximum tangential velocity at the first local peak.

Initial decision errors occurred when the initial direction of the movement was aimed toward the object on ellipse trials, or toward the bar on circle trials. Final decision errors occurred when the final hand position was closer to the object on ellipse trials or closer to the bar on circle trials. A corrected initial decision error was defined for trials in which an initial decision error occurred, but the final decision was correct.

Execution errors were identified on circle trials in which the final decision was correct, but the participant did not successfully hit the circle in the given time. Restricting the analysis to trials in which the circle was attempted to be hit allows for a comparison of No Decision blocks (in which all trials involved attempting to hit the circle), and Decision blocks (in which some trials involved an ellipse and/or a decision to avoid the object). The execution errors resulted from the cursor passing the Y position of the object without hitting it (i.e., poor trajectory), or from the hand failing to reach the Y position of the object (i.e., too slow).

### 2.5. tatistical Analyses

To compare performance and hand kinematic variables across conditions, we conducted two-way repeated measures ANOVAs using movement type (Reaching or Interception) as within-subject factor and age group (Young or Older) as between-subject factor. The alpha level for significance was set at 0.05 and effect sizes are reported using generalized η^2^. Post-hoc pairwise comparisons were conducted using the Holm correction. Linear regression was used for bivariate comparisons, with alpha level set to 0.05, and the statistical comparison of correlations between conditions was done using the Dunn and Clark’s z for dependent groups with nonoverlapping variables (Dunn and Clark, 1969), as implemented in the cocor package in R (Diedenhofen and Musch, 2015).

## 3. Results

### 3.1. Older adults show fewer corrections of initial decision errors

We first investigated decision-making performance of young and older adults based on their movement kinematics. Initial decision errors were identified on trials in which the initial hand movement direction did not match the expected movement direction (i.e., incorrectly trying to avoid a circle or hit an ellipse). Both young [*t* = −11.10, *P <* 0.001] and older adults [*t* = −8.97, *P <* 0.001] made more initial decision errors for Interception relative to Reaching [main effect of movement type: *F* (1,39) = 191.91, *P* < 0.001, *η*^*2*^ = 0.48] (Fig. 2A), suggesting that movement difficulty influenced commitment to the initial decision.

**Figure 2:**
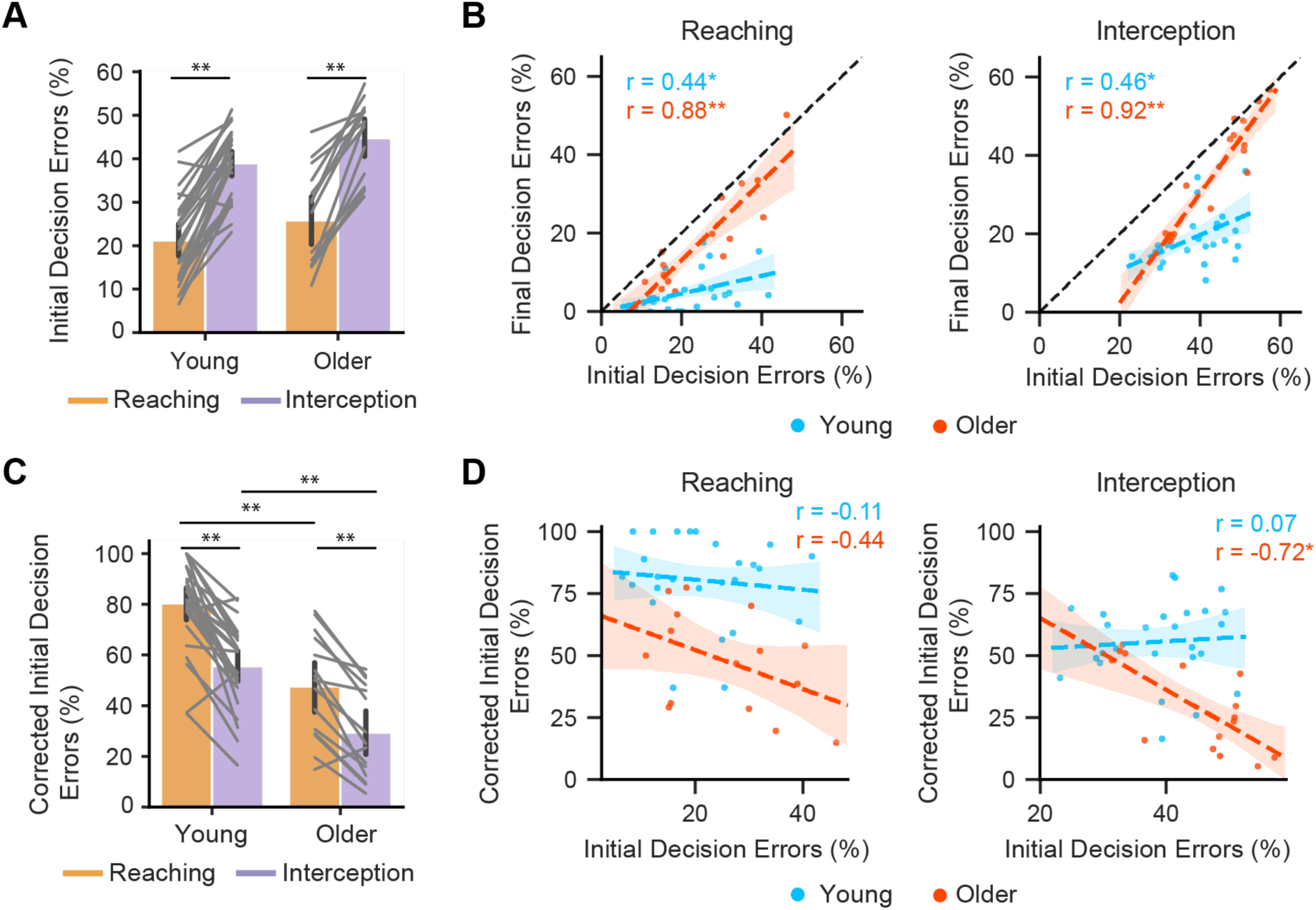
Older adults show fewer corrections of initial decision errors. A: Initial decision errors were higher during Interception for both young and older adults. B: Percentage of initial decision errors correlated more strongly with final decision errors for older adults in both Reaching and Interception. The dashed black line indicates no corrections were made during movements. C: Corrected initial decision errors (change from initial decision to final decision) occurred more frequently during Reaching than Interception. Older adults corrected fewer initial decision errors. * indicate P < 0.05, ** indicate P < 0.001.

Surprisingly, older adults did not make more initial decision errors than young adults [main effect of age: *F* (1,39) = 3.63, *P* = 0.06, *η*^*2*^ = 0.07] (Fig. 2A). However, older adults had a higher percentage of final decision errors (i.e., final position closer to bar on circle trials or to the object on ellipse trials) than younger adults [main effect of age: *F* (1,39) = 31.90, *P* < 0.001, *η*^*2*^ = 0.41], indicating that older adults were more likely to not correct their initial decision if the decision was incorrect. Indeed, though the percentage of initial and final decision errors were positively correlated for both young (Reaching: *r* = 0.44, *P* = 0.02, Interception: *r* = 0.46, *P* = 0.02) and older (Reaching: *r* = 0.88, *P* < 0.001, Interception: *r* = 0.92, *P* < 0.001) adults, the association between initial and final decisions was significantly higher for older adults for both Reaching (*z* = −2.53, *P* = 0.01) and Interception (*z* = −3.16, *P* = 0.002) (Fig. 2B).

Why were older adults less likely to correct their initial decisions? One possibility is that older adults were more constrained by the motoric demands of the task. Supporting this idea, both young [*t* = 8.30, *P* < 0.001] and older adults [*t* = 4.64, *P =* 0.001] had more corrections during Reaching than Interception [main effect of movement type: *F* (1,39) = 75.98, *P* < 0.001, *η*^*2*^ = 0.26], and young adults were much more likely to correct initial decision errors than older adults for both movement types [Reaching: *t* = −5.43, *P <* 0.001, Interception: *t* = 4.77, *P =* 0.001] [main effect of age: *F* (1,39) = 31.99, *P* < 0.001, *η*^*2*^ = 0.40] (Fig. 2C). Furthermore, older adults with more initial decision errors during Interception were also less likely to correct those errors (*r* = *-0*.*72, P* = 0.002), indicating that the initial decision errors were not simply a result of a strategy to “offload” the decision post-initiation (Fig. 2D). These results suggest that the capacity for online decision-making and movement correction is greater when the movement is easier to perform.

### 3.2. Older adults launch interception movements earlier during decision-making

As expected, the added neural processing required for judging shapes led to a significant increase in reaction time (RT) during Decision blocks relative to No Decision blocks [*t* = 22.92, *P<* 0.001]. The increase in RT for both young [*t* = 3.50, *P =* 0.004] and older adults [*t* = 9.58, *P <* 0.001] was larger for Reaching than for Interception trials [main effect of movement type: *F* (1,38) = 95.95, *P* < 0.001, *η*^*2*^ = 0.32]. Interestingly, this difference was driven mainly by the older adult group [interaction of age group and movement type: *F* (1,38) = 32.02, *P* < 0.001, *η*^*2*^ = 0.13] (Fig. 3A). For Reaching, older adults had a larger increase in RT than young adults [*t* = 2.22, *P =* 0.03]. In contrast, for Interception, older adults had a smaller increase in RT [*t* = −2.62, *P =* 0.025], implying that older adults chose to reduce decision time in order to initiate an earlier movement to intercept the object in time.

**Figure 3:**
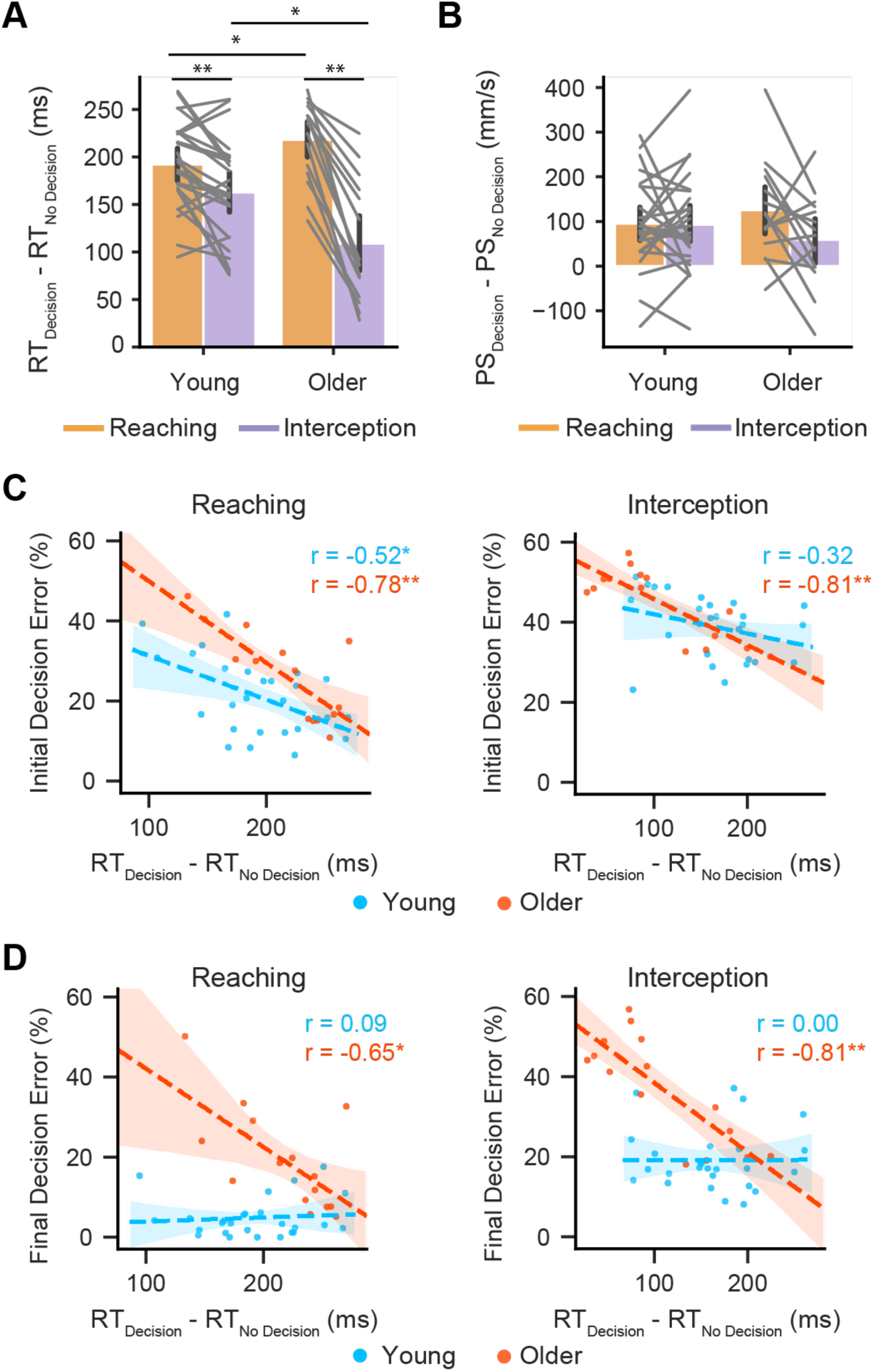
Older adults launch interception movements earlier during decision-making. A: The increase in reaction time (RT) from No Decision to Decision was larger for Reaching trials. Older adults showed a larger increase in RT during Reaching and a smaller increase in RT during Interception than young adults. B: The peak speed (PS) of the limb movement increased from the No Decision to Decision, but the increase in PS was similar across age group and movement type. C and D: The difference in reaction time from No Decision to Decision was negatively correlated with initial decision errors (C) and final decision errors (D) for older adults in both Reaching and Interception trials. * indicate P < 0.05, ** indicate P < 0.001.

We first compared movement kinematics between Decision and No Decision to eliminate any confounds in the interpretation of our results. Overall, Peak speed (PS) increased from No Decision to Decision, but there were no significant differences in the increase in PS between the two groups [main effect of age: *F* (1,39) = 0.00, *P* = 0.95, *η*^*2*^ < 0.001] or between movement types [main effect of movement type: *F* (1,39) = 3.14, *P* = 0.08, *η*^*2*^ = 0.02] (Fig. 3B). This suggests that when perceptual decision-making was added to the task, participants compensated for the longer reaction times with higher movement vigor (Summerside et al., 2018).

We then looked at how increase in RT during Decision blocks influenced initial decision errors. There was a strong negative correlation between the increase in RT to Decision from No Decision with the initial decision errors for both Reaching (*r* = −0.78, *P* < 0.001) and Interception (*r* = −0.81, *P* < 0.001) for older adults (see Fig. 3C). Furthermore, amongst older adults, the difference in RT between Decision and No Decision predicted the final performance (see Fig. 3D) in the task for both Reaching(*r* = −0.65, *P* = 0.01) and Interception (*r* = −0.81, *P* < 0.001). Thus, older adults who adjusted their RTs to be longer during Decision blocks relative to No Decision had fewer initial and final decision errors whereas older adults who “rushed” their decisions (smaller difference between Decision RT and No Decision RT) exhibited a higher number of initial and final decision errors.

For young adults, the correlation between RT adjustments and initial errors was only significant for Reaching (*r* = −0.52, *P* = 0.01) but not for Interception. Furthermore, the relationship between RT adjustments during decision-making and final decision errors was not significant in younger adults (Reaching: *r* = 0.09, *P* = 0.66; Interception: *r* = 0.00, *P* = 0.99) and significantly different from the correlations observed in older adults (Reaching: *z* = 2.45, *P* = 0.01; Interception: *z* = 3.19, *P =* 0.001). This suggests that, unlike older adults, the choice to initiate movement early was not associated with reduced decision accuracy as young adults could countermand their decision during the movement.

Overall, the results for Reaching were consistent with expectations-older adults were slower to initiate movements during Decision trials, and individuals who took longer also made fewer initial and final decision errors. In other words, older adults favored decision accuracy during reaching trials. In contrast, during Interception, older adults took 113 ± 15 ms longer in Decision blocks than the No Decision blocks, but this additional time was on average ∼50 ms shorter than the additional time taken by the young adults (163 ± 11 ms). One explanation for this is that older adults were more likely to prematurely launch the movement before they had completed the decision during Interception, resulting in more erroneous decisions.

### 3.3. Decision to move early is associated with poorer movement execution in older adults

In addition to decision errors, participants could also make errors specific to motor execution-related components of the task. These errors included poor estimate of the object’s position or an inability to adjust to the imposed time constraints. These errors were identified only on circle trials in which the final decision was correct, but participants did not successfully hit the circle in the given time.

First, our results showed that the demands of interception relative to reaching movements were greater for older adults, even in the No Decision condition. Older adults made more execution errors in Interception trials [*t* = 3.76, *P =* 0.002] than young adults [main effect of age: *F* (1,39) = 14.17, *P* < 0.001, η^2^ = 0.18, Fig. 4A]. Both young [*t* = −6.66, *P <* 0.001] and older adults [*t* = −8.07, *P <* 0.001] made more errors in Interception than Reaching trials [main effect of movement type: *F* (1,39) = 109.22, *P* < 0.001, η^2^ = 0.52]. There was a larger increase in execution errors in Interception for older adults compared to younger adults [interaction of age group and movement type: *F* (1,39) = 5.75, *P* = 0.02, *η*^*2*^ = 0.05].

**Figure 4:**
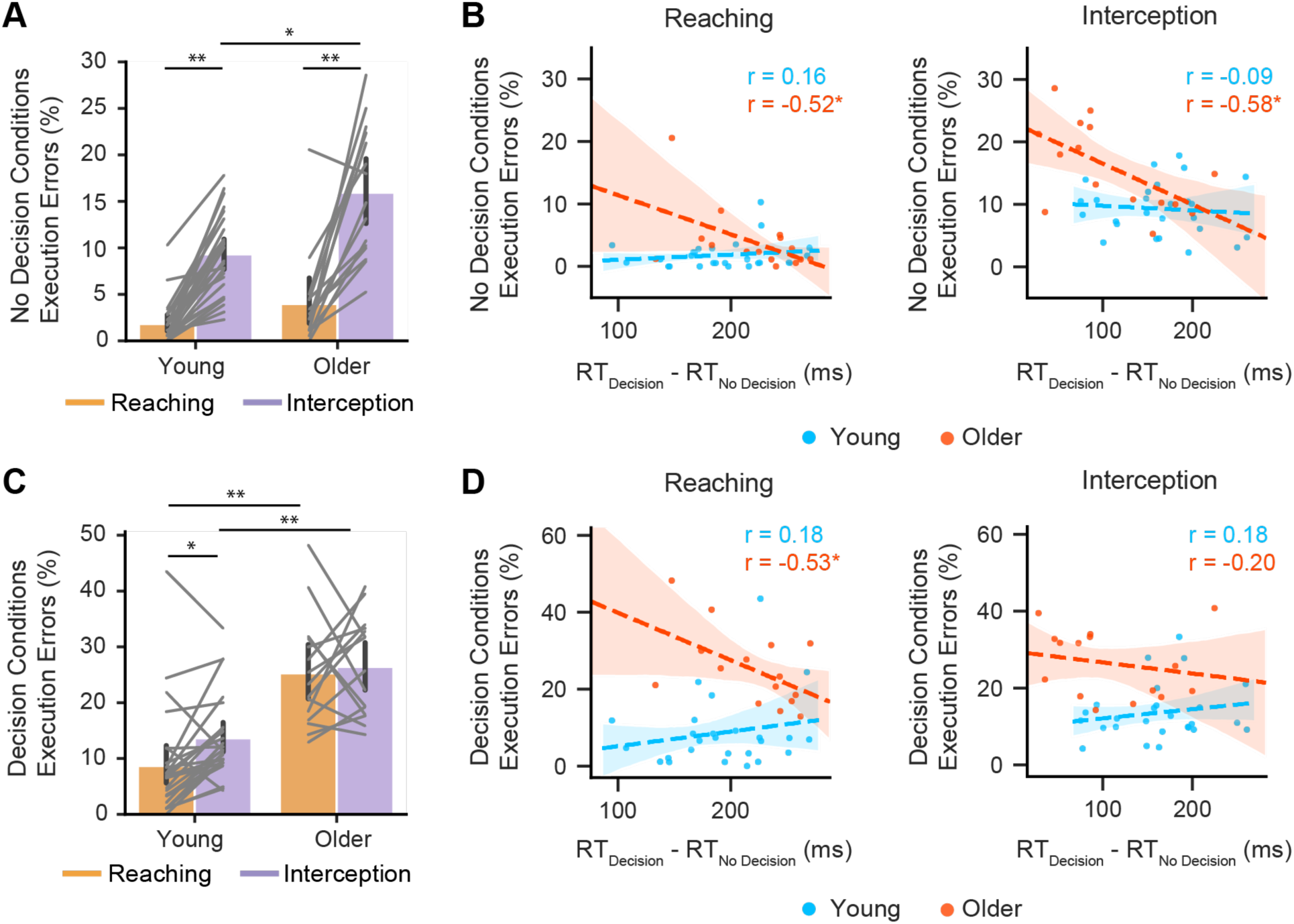
Decision to move early is associated with deficits in movement execution in older adults. A: In the No Decision conditions, there were more execution errors made during Interception trials. Older adults made more errors than younger adults. B: The difference in reaction time (RT) from No Decision to Decision was negatively correlated with execution errors for older adults in the No Decision conditions. C: In the Decision conditions, there were more execution errors made during Interception and by older adults. D: In Decision conditions, the difference in RT from No Decision to Decision was negatively correlated for older adults during Reaching. * indicate P < 0.05, ** indicate P < 0.001

We then correlated No Decision RT with the execution errors during No Decision and found no significant correlation for either young (Reaching: *r* = 0.38, *P* = 0.06; Interception: *r* = 0.22, *P* = 0.28) or older adults (Reaching: *r* = 0.19, *P* = 0.51; Interception: *r* = 0.20, *P* = 0.46). Thus, the reaction times alone were not predictive of accurate motor performance in the No Decision condition. However, the number of execution errors made during the No Decision conditions were predictive of the change in RT between Decision and No Decision conditions for both Reaching (*r* = −0.52, *P* = 0.04) and Interception (*r* = −0.58, *P* = 0.02) for older adults, but not for young adults (Reaching: *r* = 0.16, *P* = 0.43; Interception: *r =* −0.09, *P* = 0.66). These correlations were statistically different between young and older adults for Reaching (*z* = 2.09, *P* = 0.04) but not for Interception (*z* = 1.62, *P* = 0.10) (Fig. 4B). The negative correlation between these variables suggests that older adults who made more execution errors during the No Decision condition were also more likely to initiate their movements early during Decision blocks.

In the Decision blocks, older adults made more execution errors than younger adults in both Interception [*t* = 5.03, *P <* 0.001] and Reaching [*t* = 5.31, *P <* 0.001] trials [main effect of age: *F* (1,39) = 36.08, *P* < 0.001, *η*^*2*^ = 0.41, Fig. 4C]. Furthermore, the increase in execution errors during Decision (relative to No Decision) was larger for older adults [main effect of age: *F* (1,39) = 32.05, *P* < 0.001, *η*^*2*^ = 0.34] for both Reaching [*t* = 5.64, *P* < 0.001] and Interception [*t* = 3.07, *P* = 0.01]. Young adults made more errors in Interception than Reaching [*t* = −2.79, *P* = 0.02] [main effect of movement type: *F* (1,39) = 4.31, *P* = 0.04, *η*^*2*^ = 0.03] but there were no differences for older adults.

The number of execution errors in the Decision blocks were correlated with the difference in RT between Decision and No Decision for older adults in Reaching trials (*r* = −0.53, *P* = 0.04, Fig. 4D), but not for young adults (*r =* 0.18, *P* = 0.37). These correlations were also significantly different *(z* = 2.18, *P* = 0.03). Execution errors during reaching in Decision blocks were predominantly due to not reaching the object in time (hand movement was too slow)—since there was no salient cue to indicate the time constraint during Reaching, older adults may have taken more time to make their decision and consequently did not have enough time to hit the object. For Interception, the correlation between the difference in RT and execution errors was not significant for young or older adults. and there was no difference between the correlations (*z* = 1.08, *P* = 0.28).

## 4. Discussion

This study examined how aging impacts decision-making and motion-processing for visuomotor performance. To that end, young and older adults judged the shape of objects and made manual reaching and interception movements based on those decisions. We found that compared to young controls, older adults made a similar number of initial decision errors as young adults, but they corrected a smaller percentage of those errors, resulting in a lower final decision accuracy. Final decision errors were more strongly correlated with a smaller reaction time increase during decision-making in older adults than young adults, and execution errors increased during decision-making relative to young adults. Together, these results confirm our first prediction and suggest that older adults had a greater difficulty with the online adjustments necessary to successfully decide on and execute the appropriate movement. Furthermore, consistent with our second prediction, these differences were exacerbated when the task required intercepting a moving object rather than reaching a stationary object, suggesting that the capacity for online decision-making and movement corrections depends on task complexity.

### 4.1. Initial decisions made by older adults reflect a stronger commitment to an action plan

Final decision errors were typically lower than the initial decisions errors for all the participants (Fig. 2). This reflects that participants changed their mind on the initial decision during the movement. Decision-making involves accumulation of noisy evidence to produce a decision (Gold and Shadlen, 2007; Ratcliff and Smith, 2004). Previous work has shown that in a two-alternative forced choice task, participants sometimes initiate limb movements before decision-making is complete and then change their mind during the ongoing movement (Friedman et al., 2013; Resulaj et al., 2009). Resulaj and colleagues proposed a model showing that the change in the initial decision may reflect that the sensorimotor system exploits information that is still in the “processing pipeline” when the initial decision is made to subsequently either reverse or reaffirm the initial decision.

We found similar results in our study. Both older adults and young controls made similar number of initial errors, but older adults corrected fewer of those errors. Furthermore, older adults made more initial decision errors during Interception trials (Fig. 2A) and corrected a smaller percentage of those errors (Fig. 2C). This pattern of results suggest that: a) older adults may be less able to exploit sensory information in the “processing pipeline” once the limb movement is underway; and b) this capability is further diminished when they intercept moving objects. A simple interpretation of these results is that online processing of visuomotor information during movements may leave limited “bandwidth” for perceptual decision-making in older adults, minimizing the likelihood of online corrections to initial decision errors. In other words, the initial decision made by older adults is a stronger commitment to a plan of action, whereas younger adults are not fully committed to their initial decision. The motor demand required during interception movements, especially for older adults, may further reduce the likelihood for adjusting decisions post-initiation (Burk et al., 2014).

### 4.2. Early age-induced declines in dorsal stream processing may underlie execution errors

In our study participants had to make limb movements towards static (reaching) and dynamic (interception) objects. The frontoparietal areas along the dorsal visual stream are involved in the control of reaching movements (Bosco et al., 2008; Senot et al., 2008; Vesia and Crawford, 2012) through facilitation of visual attention, eye movements, motion-processing, and eye-hand coordination (Battaglia-Mayer and Caminiti, 2018; Corbetta, 1998; Medendorp et al., 2011). The different eye-hand coordination strategies required for interception movements engage additional neural areas along the dorsal stream, such as area MT+ that is involved in smooth pursuit eye movements (Delle Monache et al., 2015; Dukelow et al., 2001; Mrotek and Soechting, 2007; Spering et al., 2011). In the No Decision condition, older adults made more movement execution errors than young adults in interception movements (Fig. 4A). These deficits may be due to the early age-induced declines in dorsal stream mediated pursuit eye movements (Sharpe and Sylvester, 1978) as well as motion-processing (Langrová, J. et al., 2006; Sciberras-Lim and Lambert, 2017).

Humans produce different movement trajectories for interception movements compared to reaching movements (Smeets and Brenner, 1995). This has been attributed to a more pronounced reliance on online feedback control for interception where limb movements are regulated through continuous processing of sensory information (Lee et al., 1997; Montagne et al., 1999; Van Donkelaar et al., 1992). Online feedback control is facilitated by dorsal stream areas in the posterior parietal cortex (Day and Lyon, 2000; Pisella et al., 2000). Not surprisingly, reaching studies using the target-jump paradigm have shown deficits in online feedback control in older adults (O’Rielly and Ma-Wyatt, 2019; Sarlegna, 2006). We only measured one aspect of kinematic performance, hand peak speed (PS), and found no differences between the groups or between conditions. However, the fact that older adults made more execution errors does support the notion that online feedback control may be compromised in older adults.

### 4.3. Visuomotor decision errors suggest impaired ventral-dorsal stream processing in older adults

Shape recognition is primarily facilitated by the ventral visual stream (Breitmeyer, 2014; Konkle and Caramazza, 2013; Lehky and Sereno, 2007; Pasupathy, 2006). Accordingly, the Decision condition engaged additional areas along the ventral visual stream to differentiate the circular targets from the ellipses. Since initial decision errors were not different between older adults and young controls. This suggests that neural processing in the ventral stream may not deteriorate to the same extent as the dorsal stream, resulting in somewhat intact cognitive processing relative to deficits in motor control in older adults (Kuba et al., 2012; Ruitenberg and Koppelmans, 2020).

One interesting result in our study was that initial decision errors were higher during the interception movements for both groups (Fig. 2A). Possibly, participants in both groups initiated interception movements prematurely, before the decision-making process was complete to secure enough time to complete the movement since the total trial time was fixed and limited to 800-950 ms (Barany et al., 2020). The reaction time adjustments during Decision blocks were indeed shorter for interception movements than reaching movements for both groups, but even more pronounced among older adults (Fig. 3A). This resulted in more initial and final decision errors during interception, supporting recent evidence that decision-making is impaired when the movement required is more demanding to perform (Hesse et al., 2020; Reynaud et al., 2020). The dorsal and ventral streams are driven predominantly by magnocellular and parvocellular inputs, respectively, and axons of parvocellular cells have slower conduction velocity than magnocellular cells (Maunsell et al., 1999).

Consequently, information-processing tends to be slower in the ventral stream (Chen et al., 2007). The relative sluggishness of this pathway and the additional burden of online sensory feedback processing during interception movements may have resulted in the limb motor system initiating movements before the decisions signals in the ventral networks reached the threshold for an overt decision.

### 4.4 Subcortical areas may trigger shorter reaction times during interception movements in older adults

The reaction time (RT) adjustments made by older adults during Decision blocks of interception movements were shorter than during reaching movements. Thus, older adults launched limb movements more rapidly during Decision Interception trials. Though this seems surprising, other studies have also shown similar results where older adults made more ballistic interception movements than young adults (De Dieuleveult et al., 2018; DeGoede et al., 2001). These results suggest that older adults may have experienced an elevated sense of perceived urgency during interception movements. Though we did not observe any differences in limb kinematics (peak speed, PS) between the two groups, the shorter reaction times support this interpretation.

Another possibility is that rapidly moving stimuli may preferentially release in older adults the manual following response (Whitney et al., 2007), a short-latency and stereotyped motor response generated by direct retinotectal and tecto-reticulo-spinal pathways that target proximal arm muscles (Boehnke and Munoz, 2008; Pruszynski et al., 2010). The manual following response is a primitive protective reflex that is elicited without sufficient preparation and does not have the sophistication of voluntary motor responses. Hyperactivation of the manual following response pathway may cause an early release of these motor responses in older adults. The unsophisticated spatiotemporal characteristics of these movements may be responsible for more erroneous motor performance (Figs. 4A & C). Hyperactivation of this pathway in older adults might be caused by two factors: a) maladaptive slowing of neural processing downstream of MT+ in the parietal cortex (Justino et al., 2001); and b) dopamine depletion in the basal ganglia (Seidler et al., 2010) and consequent disinhibition of the superior colliculus. This would leave reflex pathways for the manual following response hyperexcitable in older adults (Basso et al., 1996) and cause faster and more ballistic movements, especially when the visual stimuli are moving.

## 5.0. Conclusions

In summary, our results showed that compared to young adults, older adults were less effective in correcting initial decision errors made during both reaching and interception movements. Older adults also made more decision errors and movement execution errors during interception movements than reaching movements, reflecting the role of movement complexity in online decision-making and visuomotor control. Overall, these results suggest that early age-induced declines in dorsal stream processing and the ability to incorporate ventral stream information during movement may have a strong effect on visuomotor function in older adults.

## Acknowledgements

We thank Negar Bassiri and Chris Mejias for assisting with data collection.

## Funding sources

AGG received partial support from the Universidad de Costa Rica. DAB received support from the American Heart Association (18POST34060183). A portion of this work was supported by a grant from the University of Georgia Research Foundation, Inc. to TS.

## Notes

### Competing Interest Statement

The authors have declared no competing interest.

